# Daily rhythms in lactate metabolism in the medial prefrontal cortex of mouse: Effects of light and aging

**DOI:** 10.1101/632521

**Authors:** Naomi K. Wallace, Felicity Pollard, Marina Savenkova, Ilia N. Karatsoreos

## Abstract

Aging is associated with reduced circadian (daily) rhythm amplitude in physiology and behavior, and decreased function of the prefrontal cortex (PFC). Similar effects are seen in younger mice experiencing circadian desynchrony (CD) caused by exposure to 20h light-dark cycles (T20). Given changes in PFC structure/function, underlying metabolic functioning of the PFC may also occur. We aimed to determine whether there are similarities in neurometabolism between Aged and CD mice. Using enzymatic amperometric biosensors, we recorded lactate concentration changes in the medial PFC in freely-behaving mice. Young mice displayed a circadian rhythm of lactate, which was severely blunted by CD, while Aging only changed the rhythm in constant conditions. We simultaneously probed the relationship between arousal state and PFC lactate rhythms, showing relationships between arousal state and lactate concentration, and documenting changes that occurred in CD and aging. Finally, using RT-qPCR, we found changes in genes related to metabolism and plasticity in both Aged and CD mice. Together, these data suggest both Aging and light cycle manipulation can disrupt mPFC neurometabolism.

**Highlights:**

- Lactate recordings were taken in Aged and circadian desynchronized (CD) mice.
- Lactate displayed a circadian rhythm in Control mice which was blunted in CD mice.
- The sleep state/lactate relationship was influenced by Aging, CD, and light.
- Both Aging and CD changed the expression of genes related to neurometabolism.

## 1. Introduction

Circadian (daily) rhythms are patterns of behavior and physiology that occur approximately every 24h, and are present in nearly every living organism (Roenneberg and Merrow, 2005). In mammals, these rhythms are controlled by a central brain clock residing in the suprachiasmatic nucleus of the hypothalamus (Hurd et al., 1995; Inouye and Kawamura, 1979; Lehman et al., 1984; Silver et al., 1999) which serves to synchronize the peripheral clocks found throughout the body. The maintenance of circadian rhythms and their synchrony both within an organism and between the organism and its environment, is critically important for health and well-being. Circadian desynchrony (CD) refers to a state of misalignment between circadian oscillators. This state can be induced by exposure to irregular light cycles through lifestyle factors like shift work, jet-lag, and late-night technology use, and can also arise as a consequence of normal aging (Ancoli-Israel, 2009; Monk, 2005; Spira et al., 2017).

In humans, CD is associated with many health problems including obesity (Laermans and Depoortere, 2016) and depression (Emens et al., 2009), as well as impairments in cognitive function such as decreased verbal (Rouch et al., 2005) and spatial (Cho, 2001) memory performance. Studies in rodents have further supported the detrimental effects of CD, including decreased immune function (Phillips et al., 2015), increased anxiety (Borniger et al., 2014) and reduced cognitive flexibility (Karatsoreos et al., 2011). Working memory (Hupalo and Berridge, 2016) and cognitive flexibility (Jett et al., 2017) are both associated with the prefrontal cortex (PFC), making it a good candidate brain region for exploring CD-induced neuropathology. We have previously shown that environmentally-induced CD decreases the dendritic complexity of neurons in the medial PFC (mPFC) and causes decrements in cognitive flexibility (Karatsoreos et al., 2011), demonstrating that CD can drive morphological changes in the mPFC that are associated with degraded cognitive function.

Reduced dendritic branching in the PFC (de Brabander et al., 1998), and deficits in working memory (Mattay et al., 2006) and executive function (Singh-Manoux et al., 2012) are features of aging in humans and other vertebrates. Normal aging is also accompanied by changes in circadian rhythms, including blunted temperature (Weitzman et al., 1982) and melatonin rhythms (Reiter et al., 1980), and increased sleep fragmentation (Spira et al., 2017). More severe circadian dysfunction is associated with diseases of old age, including Alzheimer’s Disease (Musiek et al., 2015) and Parkinson’s Disease (Videnovic and Golombek, 2013). Another hallmark of aging is altered neurometabolism, such as a reduction in cerebral blood flow (Fabiani et al., 2014) and changes in the concentration of various metabolites (Harris et al., 2014).

The Astrocyte Neuron Lactate Shuttle Hypothesis (ANLS) (Pellerin and Magistretti, 1994) provides a means to understand the regulation of dynamic changes in local neural energy demand. In this model, neuronal activity increases extracellular glutamate, which stimulates increased glucose uptake and glycolysis in astrocytes. This increases extracellular lactate, which can be used by nearby neurons as an energy source (Bélanger et al., 2011a). Consistent with this hypothesis, lactate increases during wake and REM sleep, when cortical firing rates are higher (Vyazovskiy et al., 2014), and decrease during NREM sleep (Naylor et al., 2011; Shram et al., 2002; Wisor et al., 2013). This leads to a 24h rhythm of lactate concentration, which is higher during the animal’s active phase than in its rest phase (Lundgaard et al., 2017; Shram et al., 2002).

Given these links between CD, neurocognitive decline, and neurometabolism we investigated whether the mPFC displays a rhythm of extracellular lactate, and if CD or aging affected this rhythm. We also investigated the relationship between lactate and sleep state in CD and advanced age. Finally, we probed the mPFC for changes in gene expression in pathways known to be involved in neurometabolic processes.

## 2. Methods

### 2.1 Mice

All mice were male C57 Bl6 mice obtained from the National Institute of Aging aged colony at Charles River Laboratories. At recording, Control and CD mice were 7 months old, and Aged mice were 19 months old. Control and Aged mice were maintained on T24, while CD mice had been in T20 for 6 weeks. All mice were single housed and had food (Purina LabDiet 5001) and water available *ad libitum* in sound attenuated and ventilated isolation cabinets (Phenome Technologies, Chicago, IL). Light at cage level was maintained at ~200 lux using white LEDs. Room temperature was maintained between 21-23°C. All mice were acclimated to our facilities for at least 1 week. All experimental procedures were approved by the Washington State University Animal Care and Use Committee.

### 2.2 RT-qPCR

Two approaches were used to detect changes in selected transcripts: Microfluidic RT-qPCR array cards or standard RT-qPCR. Mice were sacrificed at one of four time points (ZT0, 6, 12, or 18 for array cards; ZT1, 7, 13, or 19 for standard RT-qPCR) by rapid decapitation. For both experiments, there were 4-5 mice/group/time-point. For array cards, at the time of collection, PFC was isolated by rapid dissection on dry ice (Hill et al., 2010). For standard RT-qPCR, whole brains were removed and frozen in powdered dry ice and stored at −80°C until tissue punching. Tissue punches were taken using a biopsy coring tool (0.5mm ID), as previously described (Kinlein et al., 2015). In both cases, total RNA was extracted from the tissue using Direct-zol™ RNA MicroPrep Kit (Zymo Research, cat #R2060). cDNA was synthesized using the RNA samples as templates and the High-Capacity cDNA Reverse Transcription Kit with RNase Inhibitor (Life Technologies, cat. # 4374966). RT-qPCR was run with obtained cDNA using TaqMan Fast Advance Master Mix (Life Technologies, cat # 4444963), and TaqMan® Array Card 24 (Life technology, cat # 4342249, Design ID RTGZFCW) with 24 TaqMan Gene Expression Assays situated on the card and listed in Supplementary Table 1. 18s rRNA, Rn18s, and B2m were uses as housekeeping genes. Cards were run on an Applied Biosystems Viia7 RT-PCR machine. C_T_ values were produced, and raw data were analyzed using ViiA 7 RUO Software v1.2 (Applied Biosystems). Relative gene expression (Fold Change) was calculated using the comparative ΔΔC_T_ method (Schmittgen and Livak, 2008).

### 2.3 Surgical procedures

Mice were prepared for electrophysiology and lactate measurements as previously described (Clegern et al., 2012). Under isofluorane anesthesia, the skull surface was exposed, a stereotaxic apparatus was used to determine the placement of the cannula (1.95 mm anterior to Bregma, 0.25 mm lateral to the midline, −0.7 mm from skull surface), for which a hole was drilled using a 0.5mm drill bit. Four stainless steel screw electrodes were implanted, two over the frontal lobe, and two over the parietal lobe. Two additional screws were implanted above the parietal lobe to anchor the head stage. The frontal electrodes were used to measure electroencephalogram (EEG), while the parietal electrodes were used as reference and ground. These electrodes were then soldered to a printed circuit board (PCB) with a plastic 6-pin connector (Pinnacle Technology, Inc., Lawrence, KS). Two stainless steel wires attached to the PCB were inserted into the neck muscle to record electromyogram (EMG). The electrodes and PCB were then enclosed in a light-activated flowable composite resin (Prime-Dent). Mice were then allowed at least 1-week recovery before beginning recordings.

### 2.4 Extracellular Lactate and Sleep Recordings

During recording, mice were single-housed in a circular 10-inch diameter sleep chamber. Food and water were available *ad libitum*. Mice were tethered through a communicator to the sleep recording system (Pinnacle Technology, Inc.) for a 24h baseline sleep recording. During recording, mice were maintained on their previous light cycle (T20 or T24). For recordings in DD, mice were kept in their light cycle until ZT12 on day 2, after which the lights remained off. Lactate recordings began immediately after the baseline sleep recording. Lactate biosensors (Pinnacle, Inc.) were pre-calibrated with known concentrations of lactate and ascorbic acid. Approximately 1 in 5 biosensors was discarded due to ascorbic acid response. After calibration, the probes were immediately placed into the cannula of each mouse, and the mouse was re-tethered. Sleep and lactate data were acquired and archived using Sirenia Acquisition (Pinnacle Technology, Inc.). Recordings continued for ~96h, after which the biosensors were removed from their cannulas and post-calibrated. Final group sizes for these recordings were: LD Control (n=8), LD CD (n=8), LD Aged (n=9), DD Control (n=7), DD CD (n=9), DD Aged (n=10).

### 2.5 Sleep Scoring

Archived data was loaded into Sirenia Sleep Pro (Pinnacle Technology, Inc.) for analysis. Data was parsed into 10s epochs, and the power spectrum was calculated for EEG and EMG signals. Sleep scoring was done using a cluster cutting technique based on the density clusters of EEG and EMG power (Rector et al., 2009). Briefly, scatter plots were generated with EEG power on the y-axis and EMG power on the x-axis. Clusters associated with low EEG amplitude and high EMG amplitude were labeled wake, high EEG amplitude with low EMG amplitude were considered NREM, and low EEG amplitude with low EMG amplitude were considered REM. This scoring was visually confirmed based on the same criteria used to cluster score.

### 2.6 Data Analysis

Lactate data were exported from Sirenia Acquisition into Excel, trimmed, and imported to MATLAB, where outlying points were eliminated, and the downward trend was removed. For rhythm analysis, data was binned into 1h intervals. Binned data were transferred into GraphPad Prism for cosinor analysis. Time of day from the cosinor analysis was re-aligned to start at ZT18. Scored sleep files were analyzed using Sirenia’s bout analysis to determine the number and length of state bouts during light and dark periods, starting from ZT18 on Day 1 and ending ZT18 on Day 2. For the sleep-state-dependency, both lactate and sleep score were kept in 10s bins. Bouts of NREM/Wake >5 mins in the second day of recording were used. In Prism, the average lactate signal of the minute preceding the state transition was set to 100%, and the following points were expressed as a percentage of this starting value. Linear regression was used to quantify the change in lactate and to compare across groups. These data were then moved into RStudio, where they were plotted using the local regression smoothing method, with shaded areas representing SEM.

For gene expression data, Two-way ANOVAs followed by Tukey HSD post-hoc tests were used to compare differences between groups. To compare the ratio of *Ldha* to *Ldhb* transcript levels within an animal, Ct values (acquired with the same thresholding value) were subtracted from each other, and a one-tailed t-test (with Welch’s correction due to unequal variance) was undertaken to compare between day and night in each group (Young and Old). In all cases, results were considered statistically significant at the p<0.05 level.

## 3. Results

### 3.1 CD affects the daily rhythm of genes related to neurometabolism and plasticity

PFC tissue samples from Control and CD mice were taken at four time-points and gene expression was analyzed by qPCR. Twenty genes were analyzed (Table 1.). Of these, five genes had a robust rhythm in Control mice which was blunted or shifted in CD mice (Fig. 1.). Significant changes in expression pattern was found for *Gjc2* (connexin 47, Fig. 1A), *Gria2* (GluR2, Fig. 1B), *Nr3c1* (glucocorticoid receptor, Fig. 1C), *Slc1a2* (EAAT2, Fig. 1D), and *Slc2a1* (GLUT-1, Fig. 1E).

**Table 1.** Rhythmic properties of a panel of PFC transcripts that are involved in neurometabolism and plasticity, based upon delta-delta CT values. Using a cosinor analysis, amplitude and acrophase (time of highest expression) were calculated under control (LD12:12) or CD (LD10:10) conditions. N = 4/5/time point.

|  | CONTROL |  |  | CD |  |  |
| --- | --- | --- | --- | --- | --- | --- |
|  | Amplitude | Acrophase | R square | Amplitude | Acrophase | R square |
| **Cycs** | 0.1104 | 14.85 | 0.08861 | 0.1173 | 21.835 | 0.08477 |
| **Efna5** | 0.06718 | 21.04 | 0.08165 | 0.05579 | 6.969 | 0.09683 |
| **Gfap** | 0.07272 | 21.9 | 0.1201 | 0.04713 | 19.15 | 0.01517 |
| **Gjc2** | 0.1845 | 23.7099 | 0.5402 | 0.04827 | 8.664 | 0.03404 |
| **Glul** | 0.1082 | 21.125 | 0.2389 | 0.02335 | 8.884 | 0.01245 |
| **Gria2** | 0.09391 | 19.77 | 0.3394 | 0.0572 | 9.569 | 0.2266 |
| **Grin1** | 0.08769 | 20.938 | 0.231 | 0.01924 | 6.598 | 0.02348 |
| **Hk1** | 0.0614 | 21.978 | 0.1431 | 0.02335 | 2.884 | 0.01112 |
| **Ldha** | 0.1382 | 14.39 | 0.1321 | 0.1422 | 23.785 | 0.1665 |
| **Ldhb** | 0.09102 | 23.91606 | 0.1582 | 0.01709 | 19.37 | 0.005237 |
| **Nr3c1** | 0.1356 | 21.762 | 0.5526 | 0.0488 | 9.055 | 0.1363 |
| **Nr3c2** | 0.07306 | 21.972 | 0.162 | 0.06161 | 11.12 | 0.08055 |
| **Slc16a3** | 0.1569 | 23.2651 | 0.1883 | 0.05561 | 14.51 | 0.0353 |
| **Slc16a7** | 0.07 | 21.542 | 0.238 | 0.004472 | 22.23 | 0.0009785 |
| **Slc17a7** | 0.06306 | 23.0202 | 0.1238 | 0.05376 | 3.1 | 0.05598 |
| **Slc1a2** | 0.09481 | 22.247 | 0.5629 | 0.05604 | 9.678 | 0.129 |
| **Slc1a3** | 0.1179 | 21.367 | 0.534 | 0.07679 | 12.55 | 0.177 |
| **Slc2a1** | 0.1945 | 15.01 | 0.4922 | 0.05846 | 17.14 | 0.0695 |
| **Slc2a3** | 0.15 | 20.892 | 0.4963 | 0.04328 | 14.69 | 0.04089 |
| **Slco2a1** | 0.028 | 18 | 0.01059 | 0.08987 | 8.85 | 0.07384 |

**Fig. 1.**
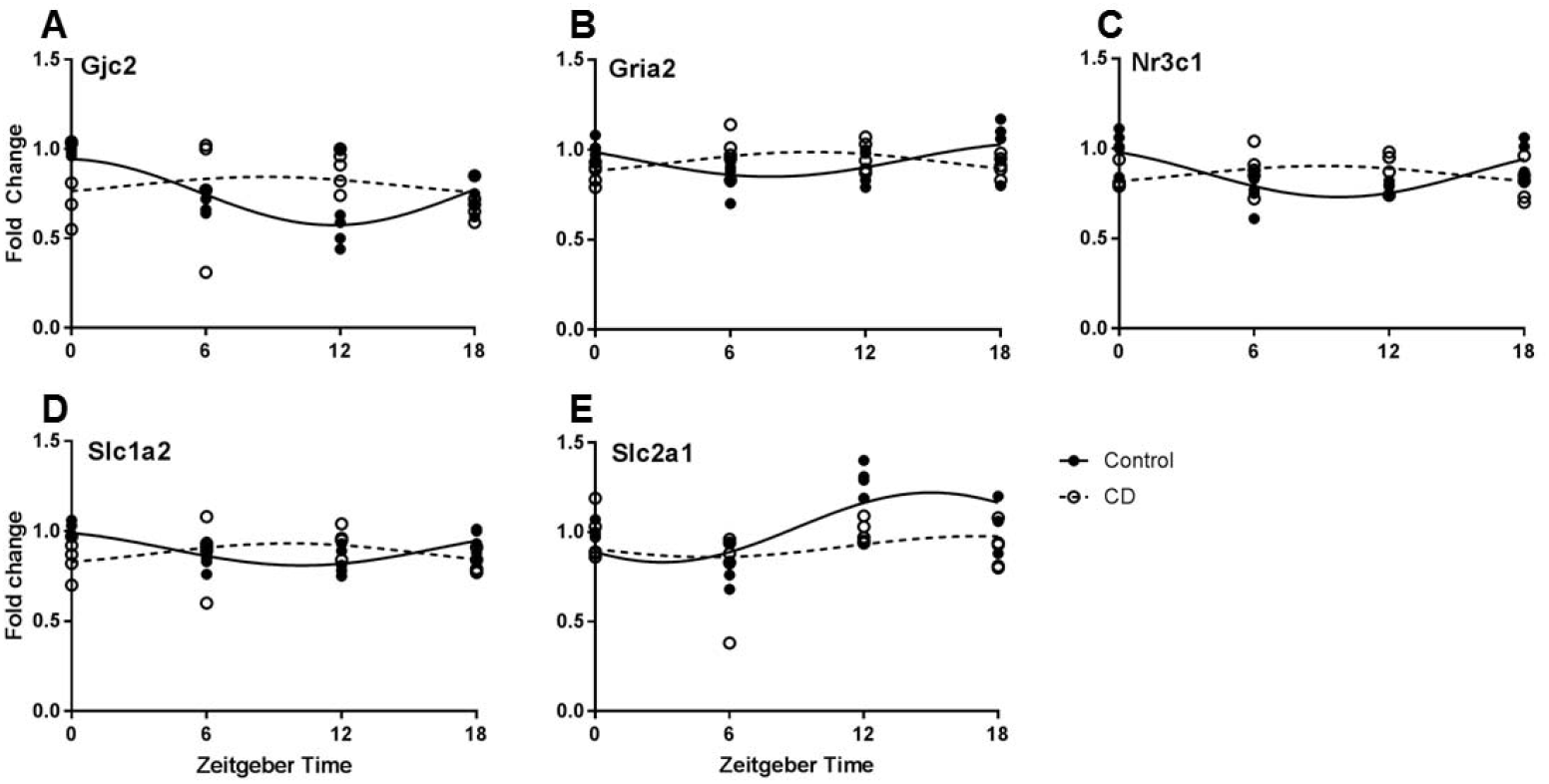
Time of day expression patterns of neurometabolic and plasticity genes is influenced by CD. **A.** For Gjc2 (connexin 47) a time of day effect was only observed in control mice, but not following CD (two-way ANOVA, interaction p=0.0374). **B.** Gria2 (GluR2) expression was highest at ZT0 and 18 in controls, but conversely was highest at ZT 6 and 12 in CD (two-way ANOVA, interaction p=0.009). **C.** Similarly, Nr3c1 (glucocorticoid receptor) expression shows highest expression at ZT0 and 18 in controls, but in CD highest expression occurs at ZT 6 and 12 (two-way ANOVA, interaction p=0.0016). **D.** The rhythm of Slc1a2 (EAAT2) expression shows a phase reversal in CD (two-way ANOVA, interaction p=0.0085). **E.** A time of day effect on Slc2a1 (GLUT-1) expression was observed in control mice, but this effect was nearly eliminated following CD (two-way ANOVA, interaction p=0.0575).

### 3.2 Sleep architecture is altered by both CD and aging

Sleep state was determined by analysis of EEG/EMG patterns. Aged mice showed shorter Wake bouts overall than Control mice (Fig. 2A). CD mice showed a reversal of the light/dark trends for sleep state. As expected, Control and Aged mice spend less time awake and more time in NREM sleep during the light phase, but CD mice did not display this difference (Fig. 2C), replicating our previous work in CD (Phillips 2015). We also analyzed NREM-Wake transitions as an indicator of sleep fragmentation, and although difference between groups were not statistically significant, there was a trend for CD mice to have more fragmented sleep (Fig. 2B). We did not find more fragmented sleep in our Aged mice.

**Fig. 2.**
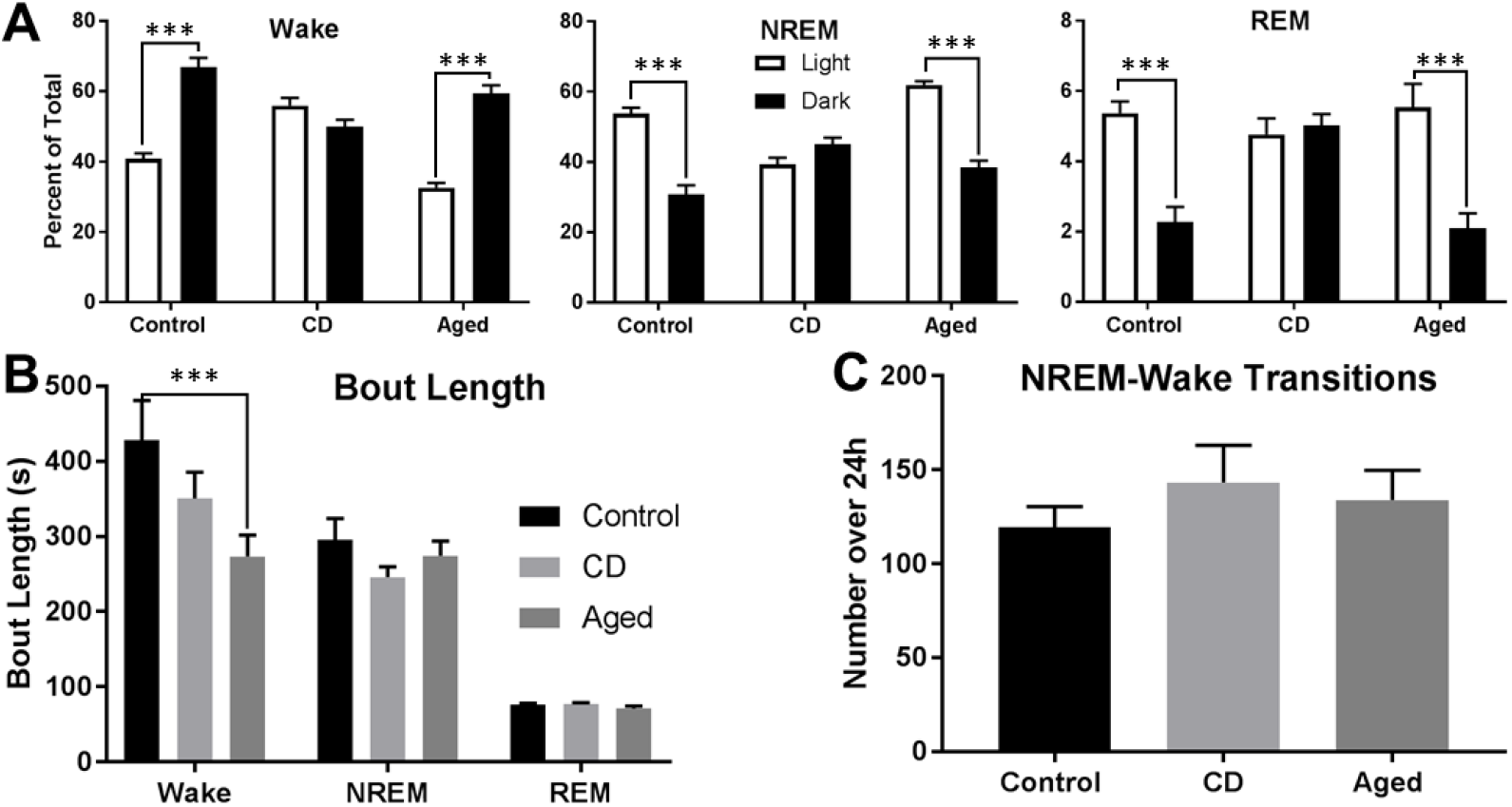
Sleep architecture is altered by both CD and aging. **A.** The average length of Wake bouts is significantly decreased in Aged mice (t-test, p=0.0002), but not in CD mice (t-test, p=0.0989). The average length of NREM and REM bouts was the same across groups.). **B.** There is a non-significant trend toward CD mice having more NREM-Wake transitions (t-test, p=0.3118), indicating increased sleep fragmentation. **C.** The relationship of sleep state and light/dark cycle is altered in CD mice. While Control and Aged mice both spend less time awake during the light phase, this difference is abolished in CD mice (t-test; Control vs. CD, p<0.0001; Aged vs. CD, p<0.0001). Control and Aged mice both spend more time in NREM and REM sleep during the light phase, and again this difference is abolished in CD mice (t-test; Control vs. CD, p<0.0001; Aged vs. CD, p<0.0001).

### 3.3 Extracellular lactate is rhythmic in the mPFC and is affected by both CD and aging

Lactate biosensors were used to provide real-time data on relative extracellular lactate concentration in freely-behaving mice. In Control mice, there was a significant diurnal rhythm of lactate in the mPFC (Fig. 3A). This rhythm was significantly blunted and phase advanced by CD (Fig. 3B), and was phase advanced by Aging (Fig. 3C). For constant darkness experiments, after recording over one light cycle, the lights went off at ZT12 and remained off for the rest of the experiment. In both Control and CD mice, constant darkness did not change the waveform of extracellular lactate (Fig. 3D, E). However, in Aged mice, the amplitude of this rhythm was significantly blunted (Fig. 3F).

**Fig. 3.**
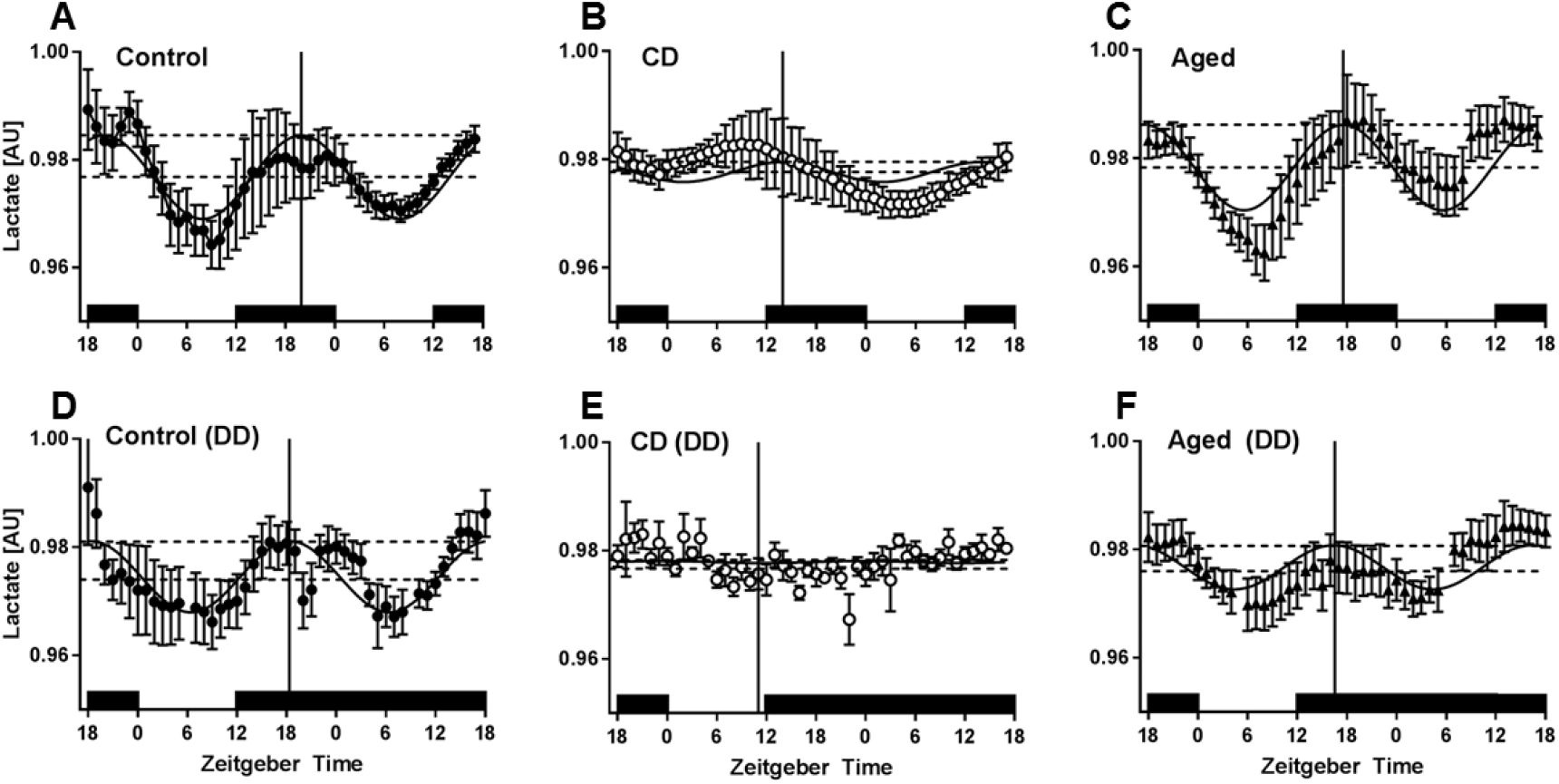
A diurnal rhythm of extracellular lactate is present in the mPFC, and this rhythm is affected by both aging and CD. **A.** Across 48h in a 12:12 LD cycle, Control mice (n=8) displayed a significant rhythm of extracellular lactate (Cosinor analysis, R^2^=0.1253, amplitude=0.007758, acrophase=26.922). **B.** CD mice (n=8) in a 10:10 LD cycle showed a different diurnal pattern of lactate (Cosinor analysis, R^2^=0.0143, amplitude=0.0007856, acrophase=20.958). Both the amplitude (t-test, p=0.0005) and the acrophase (t-test, p=0.0036) of this rhythm were significantly different from Control. **C.** Aged mice (n=9) in a 12:12 LD cycle also displayed a rhythm (Cosinor analysis, R2=0.1342, amplitude=0.007888, acrophase=24.55). While the amplitude of this rhythm was not different from that of Controls (t-test, p=0.9316) the acrophase was significantly phase advanced (t-test, p=0.0049). **D.** Mice were housed as previously, but after the first light cycle, the lights were turned off at ZT12 and did not come back on for the remainder of the experiment. Across 48h, Control mice (n=7) continued to display a significant rhythm of extracellular lactate (Cosinor analysis, R^2^=0.1214, amplitude=0.007043, acrophase=25.343). Neither the amplitude (t-test, p=0.6410) nor the acrophase (t-test, p=0.0626) were significantly different from Control mice in an LD cycle. **E.** CD mice (n=7) expressed a blunted rhythm (Cosinor analysis, R^2^=0.0003862, amplitude=0.0001919, acrophase=24.527). Neither the amplitude (t-test, p=0.1155) nor the acrophase (t-test, p=0.3606) was significantly different from CD mice in an LD cycle. **F.** Aged mice (n=10) continued to display a lactate rhythm (Cosinor analysis, R^2^=0.06399, amplitude=0.004653, acrophase=23.52). The acrophase of this rhythm was not changed by constant darkness (t-test, p=0.2407), but the amplitude was significantly blunted compared to Aged mice in an LD cycle (t-test, p=0.0245). Time of day from the cosinor analysis was re-aligned to start at ZT18. Two points were removed from the Control DD and Aged DD graphs at ZT6 on both days due to artifacts from a computer failure.

### 3.4 The relationship of sleep state and lactate is influenced by light, CD, and Aging

Scored sleep from the EEG/EMG recordings was analyzed along with lactate for bouts lasting more than 30 epochs (5 minutes). In all groups under both conditions, there was a significant decrease in lactate during NREM sleep, and a significant increase in lactate during Wake (Fig. 3). In LD, both Aging and CD blunted the lactate decrease during NREM as compared to Control mice (Fig. 3A). In DD, only CD mice showed a significantly different NREM decrease from Controls (Fig. 3B). Control and Aged mice had decreased lactate slopes during NREM in DD as compared to LD. In LD Wake bouts, only CD mice had a significantly blunted increase in lactate. In DD, only Aged mice had a significantly blunted increase. All groups showed a flattened lactate slope during Wake bouts in DD as compared to LD.

**Fig. 4.**
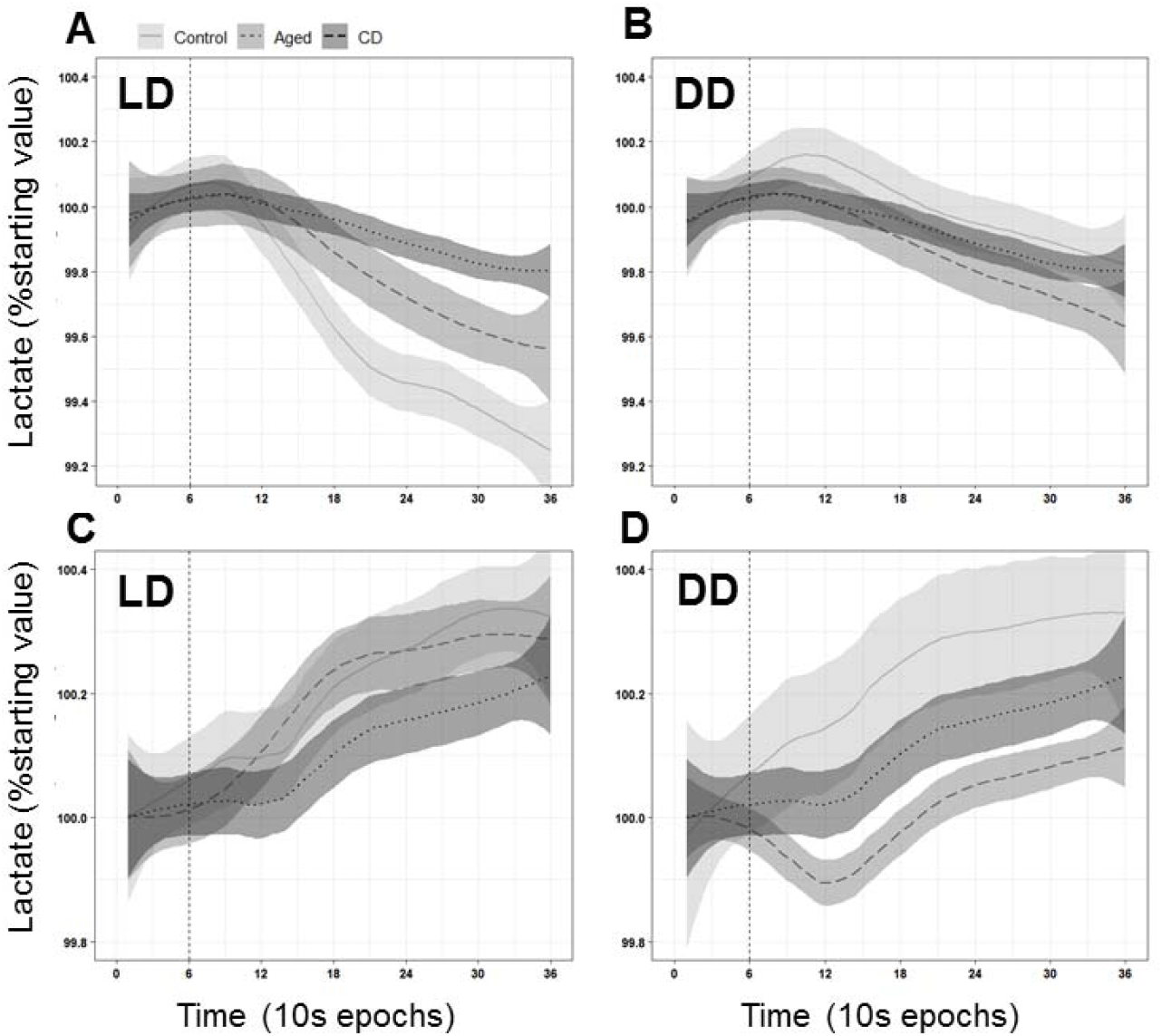
The relationship between sleep state and lactate is modified by CD, Aging, and DD. **A.** During NREM sleep, extracellular lactate levels drop in all groups (linear regression, p<0.0001). However, the slope of this decline is significantly blunted by both Aging and CD (linear regression, p<0.0001). **B.** In DD, a significant drop in extracellular lactate during NREM sleep us still observed (p<0.0001). In DD, the slope of the decline is still significantly blunted from Control by CD (p<0.0001) but not by Aging (p=0.9947). DD significantly blunted the slope of the decrease in the Control (p=0.0233) and Aged (p<0.0001) mice, but not the CD mice (p=0.5031). **C.** During Wake, extracellular lactate levels rose in all groups (Control and Aged: p<0.0001; CD: p=0.0035). This increase is significantly blunted by CD (p<0.0001), but not by Aging (p=0.1167). **D.** In DD, there continues to be a significant increase in lactate during Wake for all groups (p<0.0001). This slope was significantly blunted compared to Controls in Aged (p<0.0001) but not CD (0.6606) mice. DD significantly blunted the slope of this increase in all groups (p<0.0001).

### 3.5 Aging affects the daily expression pattern of lactate dehydrogenase enzyme mRNAs

To more specifically assess the effects of CD in the mPFC, we took fine mPFC punches at the same coordinates as the location of the biosensors (in a separate, non-surgically manipulated, set of mice). Control mice display a robust daily rhythm of *Ldha* mRNA, which codes for the enzyme that converts pyruvate to lactate. They also display a rhythm in *Ldhb*, which codes for the enzyme that converts lactate to pyruvate. This rhythm is aligned with the rhythm of extracellular lactate, such that the peak of *Ldha* expression precedes the lactate peak. In Aged mice, the rhythms in both of these enzymes are blunted.

**Fig. 5.**
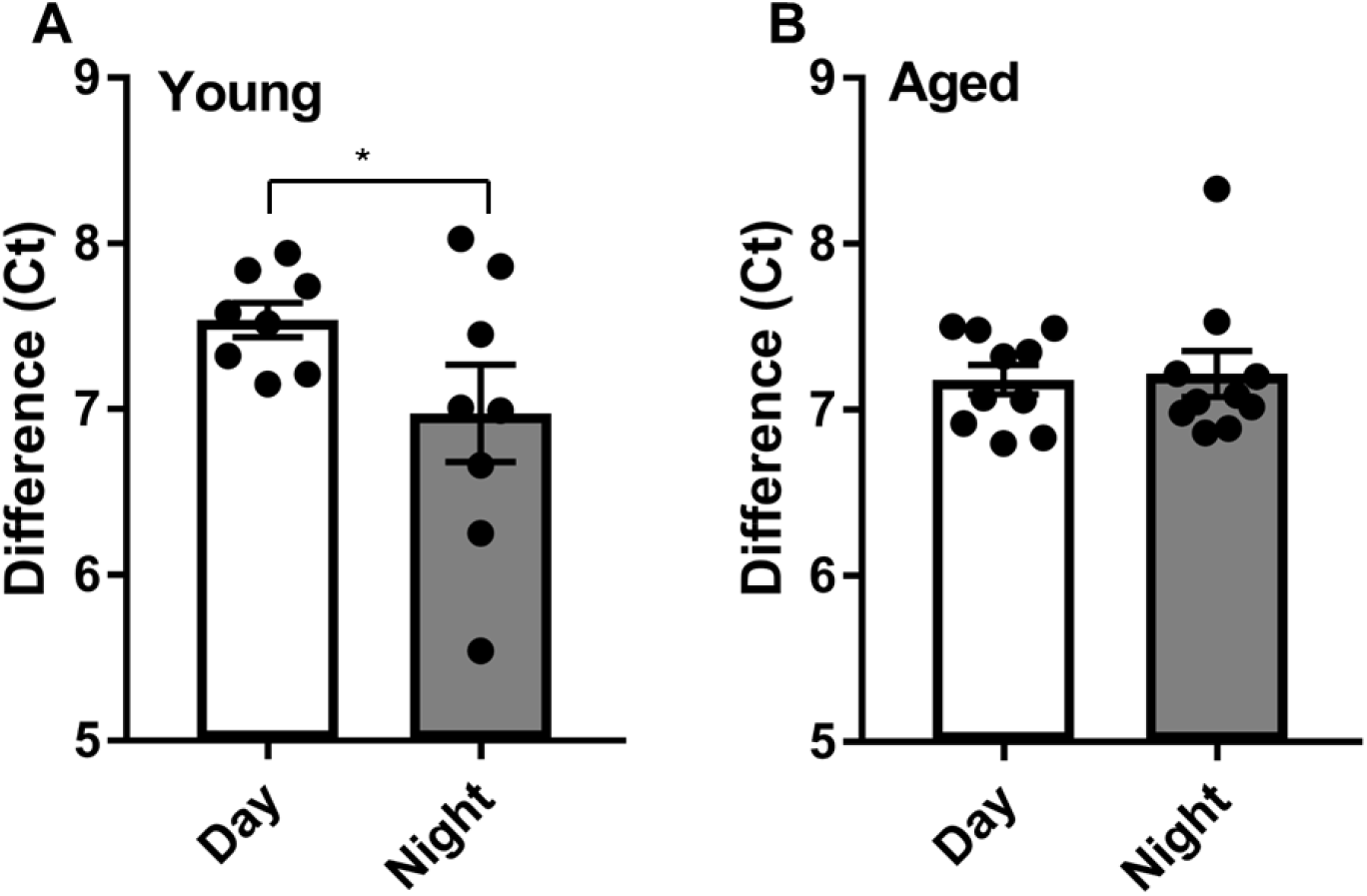
The diurnal expression of lactate dehydrogenase enzymes is blunted by aging. **A.** In Control mice, the ratio of LDHA/B changes at different times of day, with a higher ratio during the day than at night (one-tailed Welch’s t-test; p=0.05). **B.** In Aged mice, the ratio of LDHA/B does not change throughout the day (one-tailed t-test; p=0.4).

## 4. Discussion

Desynchrony of circadian rhythms affects both physical and mental health. Environmental chrono-disruption, present in shift workers, transmeridian air travelers, and most of the general public (in the form of social jet lag) is linked to metabolic disorders (Laermans and Depoortere, 2016), mental illness (Emens et al., 2009), and cognitive dysfunction (Cho, 2001; Rouch et al., 2005). The natural process of aging is also associated with changes in circadian physiology (Reiter et al., 1980; Weitzman et al., 1982), the severity of which is correlated with more pronounced health problems (Musiek et al., 2015; Videnovic and Golombek, 2013). In the current study, we examined how CD and aging affect neurometabolism of the mPFC in mice. We showed that a diurnal rhythm of extracellular lactate is present in the mouse mPFC and demonstrated that this rhythm is influenced by both CD and aging. We also replicated previous findings that demonstrated a relationship between sleep state and change in lactate concentration (Clegern et al., 2012; Naylor et al., 2011; Shram et al., 2002; Wisor et al., 2013), but further extended this to demonstrate that aging and CD can affect this relationship. With respect to gene expression changes, we found that following CD, the expression pattern of several genes involved in metabolism and neuroplasticity were altered within the mPFC. Finally, we also identified an altered expression pattern of lactate-related genes in the brain of Aged mice.

The active process by which neurons fire action potentials, and restore and maintain membrane potentials comes at a high energetic cost (Attwell and Laughlin, 2001). It is also noteworthy that neurons themselves do not keep large reserves of energetic substrates, nor the biochemical machinery that would enable rapid generation of useable energy (Cataldo and Broadwell, 1986). Thus, the energetic support for neural function likely requires complex metabolite exchange between cell types (e.g. glia). While glucose has long been considered to be the primary energy source of the brain, under some circumstances, neurons show a preference for lactate over glucose (Itoh et al., 2003), suggesting that lactate may be their primary energy substrate. This preference is somewhat puzzling, as neurons show very slow rates of glycolysis, and cannot increase this rate without compromising intracellular anti-oxidant potential (Bolaños et al., 2010). Astrocytes, on the other hand, can rapidly produce lactate, and release it into the extracellular space (Itoh et al., 2003), which has led to the articulation of the Astrocyte-Neuron Lactate Shuttle (ANLS) hypothesis (Pellerin and Magistretti, 1994). In this model, neuronal activity leads to an increase in extracellular glutamate, which stimulates glucose uptake and glycolysis in astrocytes, which release lactate into the extracellular space where it can be used by nearby neurons (Bélanger et al., 2011b). This is accomplished with a suite of proteins including lactate dehydrogenases (LDH) which convert between lactate and pyruvate (Cahn et al., 1962), the monocarboxylate transporters (MCTs) that shuttle lactate in and out of cells (Bergersen, 2007), and glutamate sensors and regulators (Bélanger et al., 2011b).

This mPFC is a key neural regulator of executive function (Hupalo and Berridge, 2016; Jett et al., 2017) and mPFC dysfunction is implicated in some mental illness (Duman and Duman, 2015; Eden et al., 2015; O’Mahony et al., 2010; Rive et al., 2013). Previous studies have demonstrated the presence of circadian rhythms in cognitive functions associated with the PFC, such as executive inhibitory control (Bratzke et al., 2012) and fear conditioning (Albrecht and Stork, 2017). Previous work in the T20 model show CD causes deficits in mPFC related behaviors such as cognitive flexibility (Jett et al., 2017; Karatsoreos et al., 2011), and a reduction in mPFC dendritic complexity (Karatsoreos et al., 2011). In the current study, we showed that there is a 24h rhythm of extracellular lactate in the mPFC. Lactate peaks during the dark phase, when nocturnal mice are most active. This finding aligns with the ANLS model’s prediction that higher levels of lactate should correspond with higher neural activity (Pellerin and Magistretti, 1994). Though a putative diurnal rhythm of lactate was previously identified in the cortex of the rat (Shram et al., 2002), these studies were somewhat limited since voltammetric recordings could only be 8-12h long. A strength of our approach is that we have demonstrated this rhythm within single individuals, continuously, over several days. We also characterized this rhythm under constant conditions to determine if the rhythm was endogenously generated (i.e. circadian). When Control mice were placed in DD, the lactate rhythm displayed no significant changes in amplitude, nor acrophase. This finding further supports our hypothesis that mPFC lactate has an endogenously generated circadian rhythm. Future studies will need to elaborate the mechanisms by which rhythm is generated, and what the relative contributions of local mPFC clocks versus the SCN master clock are.

Perhaps more significantly, we found that 6 weeks of housing in T20 significantly blunted the rhythm of lactate in the PFC. When the biological clock is disrupted, so is the lactate rhythm. Remarkably, the CD group showed no significant additional changes to their rhythm when they were placed into DD, suggesting that the T20 manipulation has fundamentally altered the endogenous rhythm of mPFC lactate.

Aged mice showed a phase advance in the mPFC lactate rhythm, relating to other work showing sleep onset and plasma cortisol levels are phase advanced in age (Yoon et al., 2003). Though aging is associated with reduced amplitude of body temperature (Weitzman et al., 1982), melatonin secretion (Reiter et al., 1980), and sleep (Spira et al., 2017), rhythms, we did not find a reduced amplitude of mPFC lactate in aged mice housed in a T24 LD cycle. However, in DD, aged mice did display a blunted lactate rhythm. Thus, our results suggest that the robust lactate rhythm observed in LD may have been a masking effect of the light cycle. Significant differences in SCN clock gene expression of aged mice has been reported, but were only apparent after housing in DD for 10 days (Nakamura et al., 2015). Since the rhythm in mPFC lactate is significantly blunted immediately upon transfer to DD, this suggests endogenous timing mechanisms within the mPFC, or synchronizing signals from the SCN to the mPFC, may be impaired in aged mice.

Our lab has previously shown that 6 weeks of exposure to T20 disrupts the circadian rhythm of sleep (Phillips et al., 2015), which was replicated in the current study. In Control mice, there is significantly more Wake and less NREM and REM sleep during the dark period than the light period. This is also true for Aged mice. However, in the CD mice, this circadian pattern of sleep state was completely abolished, no significant light/dark differences were found. When we examined average bout length, we found that Aged mice had significantly shorter bouts of Wake compared to Control mice. This finding is consistent with a previous study in 22-24 month-old mice (Wimmer et al., 2013). We also examined the NREM-Wake transitions as a measure of sleep fragmentation. A previous study from our lab found that there were significantly more NREM-Wake transitions in CD mice than Control mice (Phillips et al., 2015). In this study, the difference was not statistically significant, but the same trend was maintained.

Previous studies have identified a relationship between extracellular lactate and neuronal activity (as measured by EEG/EMG) and arousal state. Lactate levels rise during wake and REM sleep, and drop during NREM sleep (Clegern et al., 2012; Naylor et al., 2011; Shram et al., 2002; Wisor et al., 2013). This finding is well aligned with the ANLS hypothesis, which predicts that extracellular lactate levels would rise during wakefulness to support higher neural activity (Bélanger et al., 2011b; Pellerin and Magistretti, 1994). In addition, during NREM sleep or anesthesia, lactate is actively cleared from the brain by the glymphatic system (Lundgaard et al., 2017). Thus, the reduction of brain lactate is not simply a result of halting astrocytic lactate production, but instead requires active transport. This may explain why lactate levels continue to climb during sleep deprivation (Naylor et al., 2011), rather than reaching a plateau. In Control mice, we replicated and extended the findings of other groups, showing a lactate increase during wake and a decrease during NREM sleep, which was maintained in constant darkness. While this pattern was observed in all other groups, the magnitude of this change was significantly reduced by both T20 and aging. The blunted change during NREM sleep could be caused by a reduction in sleep quality. Both CD (Phillips et al., 2015) and Aged (Mander et al., 2013) mice have previously been shown to have reduced slow-wave activity during NREM sleep, supporting this idea.

To examine potential molecular correlates that might be associated with the observed effects on extracellular lactate, we examined the expression of a panel of genes related to neuroplasticity, metabolism, and transport. In relation to neuroplasticity, the expression of *Gjc2* (connexin 47) and *Gria2* (ionotropic glutamate receptor 2; GLUR2) were both altered in CD. We found that *Gjc2* was rhythmically expressed in Control mice. However, in CD mice, this time-of-day expression pattern was abolished. Connexin 47 is a key element of gap junctions between oligodendrocytes, as well as between astrocytes and oligodendrocytes, and is essential to normal myelin formation and maintenance (Fasciani et al., 2018). Mutations of *Gjc2* are known to cause a hypomyelinating leukodystrophy known as Pelizaeus-Merzbacher-like disease (Gotoh et al., 2014). Thus, it is possible that the loss of daily changes in *Gjc2* expression in CD mice could compromise the integrity of myelin or myelin dynamics. This may be particularly important if mice are exposed to CD during development, while myelination is occurring, or in situations where myelin degeneration is already occurring. The daily expression pattern of *Gria2* (GLUR2) is also affected by CD. Expression was higher at ZT 0 and 18 in Control mice, but higher at ZT12 and 16 in CD mice. GLUR2 is the subunit that renders AMPARs impermeable to Ca^2+^. After transient ischemia or epilepsy, this subunit is downregulated in vulnerable neurons before they undergo cell death (Tanaka et al., 2000). Thus, our finding may indicate that the neurons of CD mice are more vulnerable during the period when mice, and hence neurons, are the most active.

One gene broadly related to metabolism was also affected by CD: *Nr3c1* (glucocorticoid receptor; GR). Control mice showed peak expression during the night, while CD mice showed peak expression during the day. This change in expression is interesting because glucocorticoid signaling, and elements of the molecular clock are bi-directionally associated. For example, glucocorticoid signaling has been shown to be important in the synchronization of peripheral neural and body oscillators. Glucocorticoid response elements (GREs) are found in the promotor regions of several core circadian clock genes (So et al., 2009). Specifically in the PFC, it has been demonstrated that timed injections of corticosterone can modulate the timing of *Per2* mRNA expression (Woodruff et al., 2016). Conversely, the core circadian protein CLOCK is a histone acetyltransferase that represses the expression of *Nr3c1* (Liston et al., 2013). So, changes in expression of *Nr3c1* could be driven by mistiming of the molecular clock.

Finally, two solute transport genes – *Slc1a2* and *Slc2a1* – were also affected by CD. The expression of *Slc1a2* (Excitatory Amino Acid Transporter 2; EAAT2) was rhythmic in both Control and CD mice, but the peak time was reversed in CD. EAAT2 is expressed primarily on glia, where it clears glutamate from the extracellular space around synapses. A study in *Drosophila* showed that knockdown of EAAT2 increased the total amount of sleep (Stahl et al., 2018). Remarkably, our study showed that in Control mice, *Slc1a2* expression was decreased during the day; a time when mice sleep more, and thus well aligned with the genetic knockout study in flies. In CD, where the diurnal pattern of sleep is abolished, *Slc1a2* expression peaked during the night. Thus, it is possible that changes in *Slc1a2* expression and changes in sleep patterns in CD mice are related. EAAT2 acts as a trigger for lactate production in astrocytes, and there have been reports of a strong correlation between decreased EAAT2 expression and decreased lactate concentration in the hippocampus (Tsai et al., 2018). In Control mice, *Slc2a1* (Glucose Transporter 1; GLUT-1) showed a robust rhythm, with higher expression during the dark period. In CD mice, this rhythm was severely blunted. GLUT-1 is responsible for transporting glucose from blood vessels into astrocytes. Haploinsufficiency of GLUT-1 in mice reduces glucose in the cerebrospinal fluid and reduces cerebral blood flow. In yet another parallel between CD and age-related disease, defects in GLUT-1 exacerbate symptoms of Alzheimer’s disease. Mice overexpressing human amyloid β accumulate more amyloid in the brain, have increased cognitive impairment, and show greater neurodegeneration if they are also haploinsufficient for GLUT-1 (*Slc2a1*^*+/-*^*APP*^*Sw/0*^) (Winkler et al., 2015).

In our Aged mice, we also investigated the expression of two genes closely related to lactate production: *Ldha* and *Ldhb*. Lactate dehydrogenase isoform A (LDHA), is an enzyme that converts pyruvate to lactate (Cahn et al., 1962). In our study, we found a robust rhythm of *Ldha* mRNA species in Control mice, which was blunted in Aged mice. *Ldha* expression appeared to be maintained at a consistently higher level in Aged mice. The finding that *Ldha* expression is higher in Aged mice replicates the findings of an earlier study, which found elevated levels in the brains of both aged and prematurely aging transgenic mice (Ross et al., 2010). The tissue in that study was collected during the light phase (personal communication, Ross, J. M.) which corresponds with the pattern of expression that we observed, corroborating our own findings. While the previous study labeled high brain lactate as a “hallmark of aging” and implicated high *Ldha* expression as the cause, our findings add some nuance to this view. While we can neither support nor refute changes in the absolute concentration of brain lactate in old age, our findings clearly show that the expression of *Ldha* is highly time-dependent, and that older mice do not have higher expression than young mice at all times of day. *Ldhb*, on the other hand, was not modified by aging. Thus, the ratio of *Ldha* to *Ldhb* is increased during the dark period in Control mice. However, in Aged mice, this ratio remains flat throughout the day. This may explain why exposing Aged mice to constant conditions reduced the amplitude of the lactate rhythm.

### Strengths and Limitations

A strength of our approach is the use of enzyme biosensors which provide improved temporal resolution over most microdialysis protocols (Wilson and Johnson, 2008). The biosensor approach also enabled longer recordings such that we were able to record two full circadian cycles within the same individual. This is in contrast to a previous study that identified a rhythm of lactate in the rat brain, which was limited to recordings for 8-12h periods in the same individual (Shram et al., 2002). However, a limitation of our approach is that we were not able to determine absolute values of lactate, only changes in relative concentrations, meaning that we could not detect absolute differences in baseline concentration across groups.

### Summary

The work we present here supports a complex relationship between circadian rhythms, the neurometabolism of the PFC, and normal aging. We found that healthy young mice express a robust circadian rhythm of lactate neurometabolism in the mPFC, which is altered by both aging and CD. Exposure of young adult mice to a T20 cycle produces a neurometabolic phenotype that is even more severe than that of aging. Future studies can expand on this work by examining absolute lactate concentration, and by observing aged mice that have been kept in constant conditions for longer periods of time. In addition, follow-up studies should aim to elucidate the mechanisms by which this rhythm in lactate is generated within the mPFC, and the contribution of local circadian clocks in its expression. In an era of widespread light pollution, shift-work, and transmeridian travel, it is imperative that we understand the consequences of environmental circadian disruption. This study demonstrates that CD can have dire consequences for neurometabolism and contributes to the growing evidence that circadian health is essential for neurological health.

## Acknowledgements

The authors thank the vivarium staff of the Veterinary and Biomedical Research Building for their excellent animal care.

This work was supported by the National Institute of Aging (1R21AG050054-01A1) and National Science Foundation CAREER Award (1553067) to INK. A Poncin Fellowship supported NKW.

## References

Albrecht, A., Stork, O., 2017. Circadian Rhythms in Fear Conditioning: An Overview of Behavioral, Brain System, and Molecular Interactions. Neural Plast. 2017, 1–12. https://doi.org/10.1155/2017/3750307

Ancoli-Israel, S., 2009. Sleep and its disorders in aging populations. Sleep Med. 10, S7–S11. https://doi.org/10.1016/j.sleep.2009.07.004

Attwell, D., Laughlin, S.B., 2001. An Energy Budget for Signaling in the Grey Matter of the Brain. J. Cereb. Blood Flow Metab. 21, 1133–1145. https://doi.org/10.1097/00004647-200110000-00001

Bélanger, M., Allaman, I., Magistretti, P.J., 2011a. Brain Energy Metabolism: Focus on Astrocyte-Neuron Metabolic Cooperation. Cell Metab. 14, 724–738. https://doi.org/10.1016/j.cmet.2011.08.016

Bélanger, M., Allaman, I., Magistretti, P.J., 2011b. Brain Energy Metabolism: Focus on Astrocyte-Neuron Metabolic Cooperation. Cell Metab. 14, 724–738. https://doi.org/10.1016/j.cmet.2011.08.016

Bergersen, L.H., 2007. Is lactate food for neurons? Comparison of monocarboxylate transporter subtypes in brain and muscle. Neuroscience 145, 11–19. https://doi.org/10.1016/j.neuroscience.2006.11.062

Bolaños, J.P., Almeida, A., Moncada, S., 2010. Glycolysis: a bioenergetic or a survival pathway? Trends Biochem. Sci. 35, 145–149. https://doi.org/10.1016/j.tibs.2009.10.006

Borniger, J.C., McHenry, Z.D., Abi Salloum, B.A., Nelson, R.J., 2014. Exposure to dim light at night during early development increases adult anxiety-like responses. Physiol. Behav. 133, 99–106. https://doi.org/10.1016/j.physbeh.2014.05.012

Bratzke, D., Steinborn, M.B., Rolke, B., Ulrich, R., 2012. Effects of Sleep Loss and Circadian Rhythm on Executive Inhibitory Control in the Stroop and Simon Tasks. Chronobiol. Int. 29, 55–61. https://doi.org/10.3109/07420528.2011.635235

Cahn, R., Zwilling, E., Kaplan, N., Levine, L., 1962. Nature and Development of Lactic Dehydrogenases. Science 136, 962–969. https://doi.org/10.1126/science.136.3520.962

Cataldo, A.M., Broadwell, R.D., 1986. Cytochemical identification of cerebral glycogen and glucose-6-phosphatase activity under normal and experimental conditions. II. Choroid plexus and ependymal epithelia, endothelia and pericytes. J. Neurocytol. 15, 511–524.

Cho, K., 2001. Chronic’jet lag’produces temporal lobe atrophy and spatial cognitive deficits. Nat. Neurosci. 4, 567–568.

Clegern, W.C., Moore, M.E., Schmidt, M.A., Wisor, J., 2012. Simultaneous Electroencephalography, Real-time Measurement of Lactate Concentration and Optogenetic Manipulation of Neuronal Activity in the Rodent Cerebral Cortex. J. Vis. Exp. https://doi.org/10.3791/4328

de Brabander, Kramers, Uylings, 1998. Layer-specific dendritic regression of pyramidal cells with ageing in the human prefrontal cortex: Differential ageing effects in human PFC. Eur. J. Neurosci. 10, 1261–1269. https://doi.org/10.1046/j.1460-9568.1998.00137.x

Duman, C.H., Duman, R.S., 2015. Spine synapse remodeling in the pathophysiology and treatment of depression. Neurosci. Lett. 601, 20–29. https://doi.org/10.1016/j.neulet.2015.01.022

Eden, A.S., Schreiber, J., Anwander, A., Keuper, K., Laeger, I., Zwanzger, P., Zwitserlood, P., Kugel, H., Dobel, C., 2015. Emotion Regulation and Trait Anxiety Are Predicted by the Microstructure of Fibers between Amygdala and Prefrontal Cortex. J. Neurosci. 35, 6020–6027. https://doi.org/10.1523/JNEUROSCI.3659-14.2015

Emens, J., Lewy, A., Kinzie, J.M., Arntz, D., Rough, J., 2009. Circadian misalignment in major depressive disorder. Psychiatry Res. 168, 259–261. https://doi.org/10.1016/j.psychres.2009.04.009

Fabiani, M., Gordon, B.A., Maclin, E.L., Pearson, M.A., Brumback-Peltz, C.R., Low, K.A., McAuley, E., Sutton, B.P., Kramer, A.F., Gratton, G., 2014. Neurovascular coupling in normal aging: A combined optical, ERP and fMRI study. NeuroImage 85, 592–607. https://doi.org/10.1016/j.neuroimage.2013.04.113

Fasciani, I., Pluta, P., González-Nieto, D., Martínez-Montero, P., Molano, J., Paíno, C.L., Millet, O., Barrio, L.C., 2018. Directional coupling of oligodendrocyte connexin-47 and astrocyte connexin-43 gap junctions: FASCIANI ET AL. Glia 66, 2340–2352. https://doi.org/10.1002/glia.23471

Gotoh, L., Inoue, K., Helman, G., Mora, S., Maski, K., Soul, J.S., Bloom, M., Evans, S.H., Goto, Y., Caldovic, L., Hobson, G.M., Vanderver, A., 2014. GJC2 promoter mutations causing Pelizaeus–Merzbacher-like disease. Mol. Genet. Metab. 111, 393–398. https://doi.org/10.1016/j.ymgme.2013.12.001

Harris, J.L., Yeh, H.-W., Swerdlow, R.H., Choi, I.-Y., Lee, P., Brooks, W.M., 2014. High-field proton magnetic resonance spectroscopy reveals metabolic effects of normal brain aging. Neurobiol. Aging 35, 1686–1694. https://doi.org/10.1016/j.neurobiolaging.2014.01.018

Hill, M.N., Karatsoreos, I.N., Hillard, C.J., McEwen, B.S., 2010. Rapid elevations in limbic endocannabinoid content by glucocorticoid hormones in vivo. Psychoneuroendocrinology 35, 1333–1338. https://doi.org/10.1016/j.psyneuen.2010.03.005

Hupalo, S., Berridge, C.W., 2016. Working Memory Impairing Actions of Corticotropin-Releasing Factor (CRF) Neurotransmission in the Prefrontal Cortex. Neuropsychopharmacology 41, 2733.

Hurd, M.W., Zimmer, K.A., Lehman, M.N., Ralph, M.R., 1995. Circadian locomotor rhythms in aged hamsters following suprachiasmatic transplant. Am. J. Physiol.-Regul. Integr. Comp. Physiol. 269, R958–R968.

Inouye, S.T., Kawamura, H., 1979. Persistence of circadian rhythmicity in a mammalian hypothalamic “island” containing the suprachiasmatic nucleus. Proc. Natl. Acad. Sci. 76, 5962–5966. https://doi.org/10.1073/pnas.76.11.5962

Itoh, Y., Esaki, T., Shimoji, K., Cook, M., Law, M.J., Kaufman, E., Sokoloff, L., 2003. Dichloroacetate effects on glucose and lactate oxidation by neurons and astroglia in vitro and on glucose utilization by brain in vivo. Proc. Natl. Acad. Sci. 100, 4879–4884.

Jett, J.D., Bulin, S.E., Hatherall, L.C., McCartney, C.M., Morilak, D.A., 2017. Deficits in cognitive flexibility induced by chronic unpredictable stress are associated with impaired glutamate neurotransmission in the rat medial prefrontal cortex. Neuroscience 346, 284–297. https://doi.org/10.1016/j.neuroscience.2017.01.017

Karatsoreos, I.N., Bhagat, S., Bloss, E.B., Morrison, J.H., McEwen, B.S., 2011. Disruption of circadian clocks has ramifications for metabolism, brain, and behavior. Proc. Natl. Acad. Sci. 108, 1657–1662. https://doi.org/10.1073/pnas.1018375108

Laermans, J., Depoortere, I., 2016. Chronobesity: role of the circadian system in the obesity epidemic: Role of circadian clocks in obesity. Obes. Rev. 17, 108–125. https://doi.org/10.1111/obr.12351

Lehman, M.N., Bittman, E.L., Newman, S.W., 1984. Role of the hypothalamic paraventricular nucleus in neuroendocrine responses to daylength in the golden hamster. Brain Res. 308, 25–32. https://doi.org/10.1016/0006-8993(84)90913-2

Liston, C., Cichon, J.M., Jeanneteau, F., Jia, Z., Chao, M.V., Gan, W.-B., 2013. Circadian glucocorticoid oscillations promote learning-dependent synapse formation and maintenance. Nat. Neurosci. 16, 698–705. https://doi.org/10.1038/nn.3387

Lundgaard, I., Lu, M.L., Yang, E., Peng, W., Mestre, H., Hitomi, E., Deane, R., Nedergaard, M., 2017. Glymphatic clearance controls state-dependent changes in brain lactate concentration. J. Cereb. Blood Flow Metab. 37, 2112–2124. https://doi.org/10.1177/0271678X16661202

Mander, B.A., Rao, V., Lu, B., Saletin, J.M., Lindquist, J.R., Ancoli-Israel, S., Jagust, W., Walker, M.P., 2013. Prefrontal atrophy, disrupted NREM slow waves and impaired hippocampal-dependent memory in aging. Nat. Neurosci. 16, 357–364. https://doi.org/10.1038/nn.3324

Mattay, Venkata.S., Fera, F., Tessitore, A., Hariri, A.R., Berman, K.F., Das, S., Meyer-Lindenberg, A., Goldberg, T.E., Callicott, J.H., Weinberger, D.R., 2006. Neurophysiological correlates of age-related changes in working memory capacity. Neurosci. Lett. 392, 32–37. https://doi.org/10.1016/j.neulet.2005.09.025

Monk, T.H., 2005. Aging human circadian rhythms: conventional wisdom may not always be right. J. Biol. Rhythms 20, 366–374.

Musiek, E.S., Xiong, D.D., Holtzman, D.M., 2015. Sleep, circadian rhythms, and the pathogenesis of Alzheimer Disease. Exp. Mol. Med. 47, e148. https://doi.org/10.1038/emm.2014.121

Nakamura, T.J., Nakamura, W., Tokuda, I.T., Ishikawa, T., Kudo, T., Colwell, C.S., Block, G.D., 2015. Age-Related Changes in the Circadian System Unmasked by Constant Conditions. eNeuro 2. https://doi.org/10.1523/ENEURO.0064-15.2015

Naylor, E., Aillon, D.V., Gabbert, S., Harmon, H., Johnson, D.A., Wilson, G.S., Petillo, P.A., 2011. Simultaneous real-time measurement of EEG/EMG and l-glutamate in mice: A biosensor study of neuronal activity during sleep. J. Electroanal. Chem. 656, 106–113. https://doi.org/10.1016/j.jelechem.2010.12.031

O’Mahony, C.M., Sweeney, F.F., Daly, E., Dinan, T.G., Cryan, J.F., 2010. Restraint stress-induced brain activation patterns in two strains of mice differing in their anxiety behaviour. Behav. Brain Res. 213, 148–154. https://doi.org/10.1016/j.bbr.2010.04.038

Pellerin, L., Magistretti, P.J., 1994. Glutamate uptake into astrocytes stimulates aerobic glycolysis: a mechanism coupling neuronal activity to glucose utilization. Proc. Natl. Acad. Sci. 91, 10625–10629.

Phillips, D.J., Savenkova, M.I., Karatsoreos, I.N., 2015. Environmental disruption of the circadian clock leads to altered sleep and immune responses in mouse. Brain. Behav. Immun. 47, 14–23. https://doi.org/10.1016/j.bbi.2014.12.008

Rector, D.M., Schei, J.L., Rojas, M.J., 2009. Mechanisms underlying state dependent surface-evoked response patterns. Neuroscience 159, 115–126. https://doi.org/10.1016/j.neuroscience.2008.11.031

Reiter, R.J., Richardson, B.A., Johnson, L.Y., Ferguson, B.N., Dinh, D.T., 1980. Pineal Melatonin Rhythm: Reduction in Aging Syrian Hamsters. Sci. New Ser. 210, 1372–1373.

Rive, M.M., van Rooijen, G., Veltman, D.J., Phillips, M.L., Schene, A.H., Ruhé, H.G., 2013. Neural correlates of dysfunctional emotion regulation in major depressive disorder. A systematic review of neuroimaging studies. Neurosci. Biobehav. Rev. 37, 2529–2553. https://doi.org/10.1016/j.neubiorev.2013.07.018

Roenneberg, T., Merrow, M., 2005. Circadian clocks -- the fall and rise of physiology. Nat. Rev. Mol. Cell Biol. 6, 965–971. https://doi.org/10.1038/nrm1766

Ross, J.M., Öberg, J., Brené, S., Coppotelli, G., Terzioglu, M., Pernold, K., Goiny, M., Sitnikov, R., Kehr, J., Trifunovic, A., others, 2010. High brain lactate is a hallmark of aging and caused by a shift in the lactate dehydrogenase A/B ratio. Proc. Natl. Acad. Sci. 107, 20087–20092.

Rouch, I., Wild, P., Ansiau, D., Marquié, J.-C., 2005. Shiftwork experience, age and cognitive performance. Ergonomics 48, 1282–1293. https://doi.org/10.1080/00140130500241670

Schmittgen, T., Livak, K., 2008. Analyzing real-time PCR data by the comparative CT method. Nat. Protoc. 3, 1101–1108.

Shram, N., Netchiporouk, L., Cespuglio, R., 2002. Lactate in the brain of the freely moving rat: voltammetric monitoring of the changes related to the sleep-wake states: Sleep-waking and brain extracellular lactate. Eur. J. Neurosci. 16, 461–466. https://doi.org/10.1046/j.1460-9568.2002.02081.x

Silver, R., Sookhoo, A.I., LeSauter, J., Stevens, P., Jansen, H.T., Lehman, M.N., 1999. Multiple regulatory elements result in regional specificity in circadian rhythms of neuropeptide expression in mouse SCN. Neuroreport 10, 3165–3174.

Singh-Manoux, A., Kivimaki, M., Glymour, M.M., Elbaz, A., Berr, C., Ebmeier, K.P., Ferrie, J.E., Dugravot, A., 2012. Timing of onset of cognitive decline: results from Whitehall II prospective cohort study. BMJ 344, d7622–d7622. https://doi.org/10.1136/bmj.d7622

So, A.Y.-L., Bernal, T.U., Pillsbury, M.L., Yamamoto, K.R., Feldman, B.J., 2009. Glucocorticoid regulation of the circadian clock modulates glucose homeostasis. Proc. Natl. Acad. Sci. 106, 17582–17587. https://doi.org/10.1073/pnas.0909733106

Spira, A.P., Stone, K.L., Redline, S., Ensrud, K.E., Ancoli-Israel, S., Cauley, J.A., Yaffe, K., 2017. Actigraphic Sleep Duration and Fragmentation in Older Women: Associations With Performance Across Cognitive Domains. Sleep 40. https://doi.org/10.1093/sleep/zsx073

Stahl, B.A., Peco, E., Davla, S., Murakami, K., Caicedo Moreno, N.A., van Meyel, D.J., Keene, A.C., 2018. The Taurine Transporter Eaat2 Functions in Ensheathing Glia to Modulate Sleep and Metabolic Rate. Curr. Biol. 28, 3700–3708.e4. https://doi.org/10.1016/j.cub.2018.10.039

Tanaka, H., Grooms, S.Y., Bennett, M.V.L., Zukin, R.S., 2000. The AMPAR subunit GluR2: still front and center-stage. Brain Res. 18.

Tsai, S.-F., Chen, Y.-W., Kuo, Y.-M., 2018. High-fat diet reduces the hippocampal content level of lactate which is correlated with the expression of glial glutamate transporters. Neurosci. Lett. 662, 142–146. https://doi.org/10.1016/j.neulet.2017.10.024

Videnovic, A., Golombek, D., 2013. Circadian and sleep disorders in Parkinson’s disease. Exp. Neurol. 243, 45–56. https://doi.org/10.1016/j.expneurol.2012.08.018

Vyazovskiy, V.V., Cui, N., Rodriguez, A.V., Funk, C., Cirelli, C., Tononi, G., 2014. The Dynamics of Cortical Neuronal Activity in the First Minutes after Spontaneous Awakening in Rats and Mice. Sleep 37, 1337–1347. https://doi.org/10.5665/sleep.3926

Weitzman, E.D., Moline, M.L., Czeisler, C.A., Zimmerman, J.C., 1982. Chronobiology of aging: Temperature, sleep-wake rhythms and entrainment. Neurobiol. Aging 3, 299–309. https://doi.org/10.1016/0197-4580(82)90018-5

Wilson, G.S., Johnson, M.A., 2008. In-Vivo Electrochemistry: What Can We Learn about Living Systems? Chem. Rev. 108, 2462–2481. https://doi.org/10.1021/cr068082i

Wimmer, M.E., Rising, J., Galante, R.J., Wyner, A., Pack, A.I., Abel, T., 2013. Aging in Mice Reduces the Ability to Sustain Sleep/Wake States. PLoS ONE 8, e81880. https://doi.org/10.1371/journal.pone.0081880

Winkler, E.A., Nishida, Y., Sagare, A.P., Rege, S.V., Bell, R.D., Perlmutter, D., Sengillo, J.D., Hillman, S., Kong, P., Nelson, A.R., Sullivan, J.S., Zhao, Z., Meiselman, H.J., Wenby, R.B., Soto, J., Abel, E.D., Makshanoff, J., Zuniga, E., De Vivo, D.C., Zlokovic, B.V., 2015. GLUT1 reductions exacerbate Alzheimer’s disease vasculo-neuronal dysfunction and degeneration. Nat. Neurosci. 18, 521–530. https://doi.org/10.1038/nn.3966

Wisor, J.P., Rempe, M.J., Schmidt, M.A., Moore, M.E., Clegern, W.C., 2013. Sleep Slow-Wave Activity Regulates Cerebral Glycolytic Metabolism. Cereb. Cortex 23, 1978–1987. https://doi.org/10.1093/cercor/bhs189

Woodruff, E.R., Chun, L.E., Hinds, L.R., Spencer, R.L., 2016. Diurnal Corticosterone Presence and Phase Modulate Clock Gene Expression in the Male Rat Prefrontal Cortex. Endocrinology 157, 1522–1534. https://doi.org/10.1210/en.2015-1884

Yoon, I.-Y., Kripke, D.F., Elliott, J.A., Youngstedt, S.D., Rex, K.M., Hauger, R.L., 2003. Age-Related Changes of Circadian Rhythms and Sleep-Wake Cycles. J. Am. Geriatr. Soc. 51, 1085–1091. https://doi.org/10.1046/j.1532-5415.2003.51356.x

