## Supplemental Table 1 for "Daily rhythms in lactate metabolism in the medial prefrontal cortex of mouse: Effects of light and aging"

**Supplemental Information**

Supplemental Table 1

| Gene Symbol | TaqMan Assay ID |
| --- | --- |
| Aqp4 | Mm00802131_m1 |
| Cycs | Mm01621044_g1 |
| Efna5 | Mm01237700_m1 |
| Gfap | Mm01253033_m1 |
| Gjc2 | Mm04212188_m1 |
| Glul | Mm00725701_s1 |
| Gria2 | Mm00442822_m1 |
| Grin1 | Mm00433790_m1 |
| Hk1 | Mm00439344_m1 |
| Ldha | Mm00495282_g1 |
| 18s rRNA* | Hs99999901_s1 |
| Rn18s* | Mm03928990_g1 |
| B2m* | Mm00437762_m1 |
| Ldhb | Mm00493146_m1 |
| Nr3c1 | Mm00433832_m1 |
| Nr3c2 | Mm01241596_m1 |
| Slc16a3 | Mm00446102_m1 |
| Slc16a7 | Mm00441442_m1 |
| Slc17a7 | Mm00812886_m1 |
| Slc1a2 | Mm01275814_m1 |
| Slc1a3 | Mm00600697_m1 |
| Slc2a1 | Mm00441480_m1 |
| Slc2a3 | Mm00441483_m1 |
| Slco2a1B | Mm00459638_m1 |

**Table S1.** Reference for the TaqMan Assay IDs for RT-qPCR experiment. * indicates housekeeping genes.
